# Mapping Human Hematopoietic Hierarchy at Single Cell Resolution by Microwell-seq

**DOI:** 10.1101/127217

**Authors:** Shujing Lai, Yang Xu, Wentao Huang, Mengmeng Jiang, Haide Chen, Fang Ye, Renying Wang, Yunfei Qiu, Xinyi Jiang, Daosheng Huang, Jie Mao, Yanwei Li, Yingru Lu, Jin Xie, Qun Fang, Tiefeng Li, He Huang, Xiaoping Han, Guoji Guo

## Abstract

The classical hematopoietic hierarchy, which is mainly built with fluorescence-activated cell sorting (FACS) technology, proves to be inaccurate in recent studies. Single cell RNA-seq (scRNA-seq) analysis provides a solution to overcome the limit of FACS-based cell type definition system for the dissection of complex cellular hierarchy. However, large-scale scRNA-seq is constrained by the throughput and cost of traditional methods. Here, we developed Microwell-seq, a high-throughput and low-cost scRNA-seq platform using extremely simple devices. Using Microwell-seq, we constructed a single-cell resolution transcriptome atlas of human hematopoietic differentiation hierarchy by profiling more than 50,000 single cells throughout adult human hematopoietic system. We found that adult human hematopoietic stem and progenitor cell (HSPC) compartment is dominated by progenitors primed with lineage specific regulators. Our analysis revealed differentiation pathways for each cell types, through which HSPCs directly progress to lineage biased progenitors before differentiation. We propose a revised adult human hematopoietic hierarchy independent of oligopotent progenitors. Our study also demonstrates the broad applicability of Microwell-seq technology.

## Introduction

The classical model of hematopoiesis is a branched tree, rooted from long-term hematopoietic stem cell (LT-HSC) and followed by multipotent, oligopotent and unipotent progenitor stages (Akashi et al., 2000; Doulatov et al., 2012; Kondo et al., 1997; Orkin and Zon, 2008). However, this model is mainly based on the FACS cell type definition system, which is heavily biased due to the limited choices of surface markers. Several studies using *in vivo* tracing challenged this classical model and showed the hematopoiesis were actually maintained by large kinds of progenitors which each have strong lineage bias (Busch et al., 2015; Sun et al., 2014). Single cell qPCR using hundreds of marker genes also demonstrates the strong heterogeneity within the hematopoietic progenitor population (Guo et al., 2013). Recently, by coupling single cell qPCR assay and single cell differentiation assay, Notta et al. showed that Er-Mk lineages emerges directly from multipotent cells, and proposed a revised human hematopoietic hierarchy (Notta et al., 2016). However, the selection of cell populations from these studies still depends on classical FACS gating scheme. A systematic single cell atlas of human hematopoietic transcriptome with minimal influence from traditional hierarchy is lacking.

scRNA-seq is a powerful technology for profiling the whole transcriptome of individual cells (Hashimshony et al., 2012; Ramskold et al., 2012; Tang et al., 2009; Treutlein et al., 2014), which can be used to dissect cellular heterogeneity and construct lineage hierarchy. However, traditional scRNA-seq methods are expensive and time-consuming for large-scale experiments. Thus, recent studies using traditional scRNA-seq to investigate the heterogeneity of human HSPCs are not comprehensive due to their scant cell quantity (Notta et al., 2016; Velten et al., 2017). Lately, several high-throughput scRNA-seq methods have been developed, including Drop-seq and inDrop (Klein et al., 2015; Macosko et al., 2015). Nevertheless, Dropseq and inDrop heavily rely on sophisticated microfluidic devices and pump systems, which constrain their extensive application in small labs. Here, we report Microwell-seq, an extremely simple method utilizing an agrose-made microwell array and barcoded beads to profile thousands of single cells for transcriptome analysis. Microwell-seq demonstrates great advantages in convenience and simplicity, which makes it accessible to every lab.

Using Microwell-seq, we constructed a transcriptomic atlas of human hematopoietic hierarchy by profiling the transcriptome of more than 50,000 single cells, which is by far the most comprehensive atlas of *in vivo* human HSPC population. The atlas showed that adult human HSPC compartment is dominated by progenitors in a lineage-primed status while the classical oligopotent HSPCs are not observed. Unique transcriptional priming pathways can be reconstructed by combining the differentiated CD34^−^ cells. In further functional assay experiments, the lineage-primed HSPCs showed strong commitment preference to specific cell type, which validated our finding in the transcriptomic atlas. Thus, we propose a revised adult human hematopoietic hierarchy in which HSPCs directly progress to unipotent progenitors without the oligopotent stage, and the pool of unipotent progenitors will give rise to the various kinds of blood cells.

## Results

### Microwell-seq is a convenient, low-cost and simple platform for high-throughput scRNA-seq

In Microwell-seq, an agarose microarray is used to trap individual cells; and magnetic beads are used to capture the mRNAs. The diagrams showing the process of bead synthesis and microarray fabrication are shown in Figure 1. Both procedures are extremely simple. The barcoded beads are synthesized by 3 rounds split-pool (Figure 1A). Every oligonucleotide consists of a primer sequence, a cell barcode, a unique molecular identifiers (UMI) and a poly T tail (Fan et al., 2015; Islam et al., 2014; Klein et al., 2015; Macosko et al., 2015). In the first-round split-pool, magnetic beads coated with carboxyl groups are randomly distributed into a 96-well plate where the 5’ amino modified oligonucleotides are conjugated to the beads. The oligonucleotides in each well have a unique barcode sequence. Then the beads are pooled and split into another 96-well plate where the second barcode sequence is introduced by single cycle PCR. In the final-round split-pool, the third barcoded sequence, unique molecular identifier (UMI) (Islam et al., 2014) and PolyT tail are introduced (Table S1). After the split-pool, all the oligonucleotides on the same bead will have the same cell barcode and different UMI, while oligonucleotides on different beads will have different cell barcodes. The cell barcodes will be used to demultiplex the pooled sequencing results to individual cells. The UMI will be used to track single molecules to reduce the quantitative bias introduced during amplification. This beads synthesis process is easy to perform for all labs.

**Figure 1.**
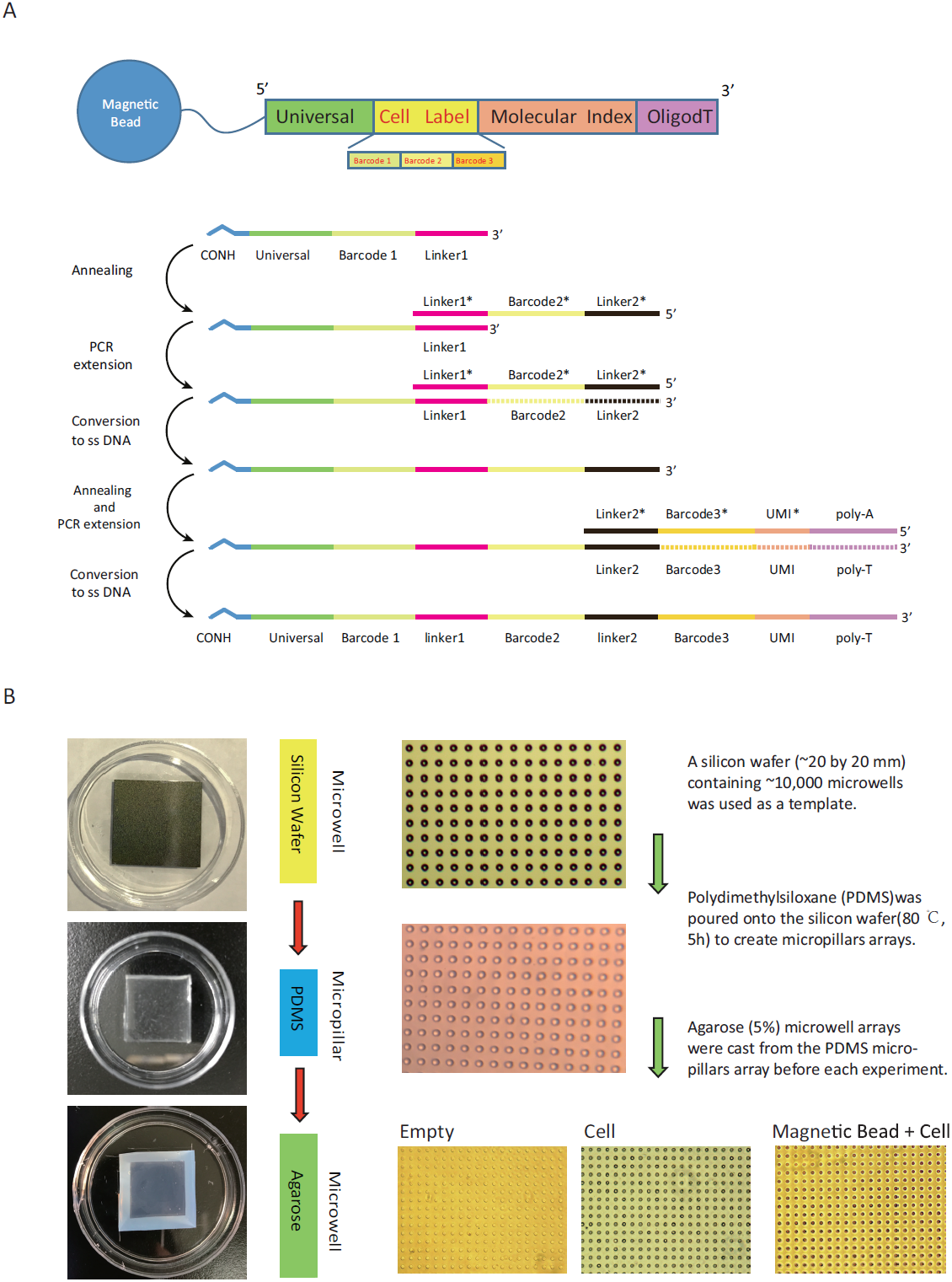
Microarray fabrication and beads synthesis. (A) The synthesis of barcoded beads. Three rounds of split-pool are used to introduce 3 parts of the oligonucleotides on the beads. After the final split-pool, all oligonucleotides on the same beads will have same cell barcode and different UMIs, and different beads will have different cell barcodes. (B) The fabrication of microarray. The microarray fabrication follows a strategy using a silicon microarray chip to make micropillar PDMS chip and then agarose microarray chip. Both silicon and PDMS chip are reusable, which greatly reduces cost and saves time. See also Figure S1 and Table S1.

The fabrication of microarray is also very simple and cheap (Figure 1B). First, a silicon chip with 100,000 microwells on the surface is manufactured and can be used as a permanent mold. Second, a PDMS chip with 100,000 micropillars is made by simply pouring PDMS on the surface of silicon chip. Finally, an agarose microwell chip is made by pouring agarose to the PDMS chip. The advantage of this method is that both silicon chip and PDMS chip are reusable, which means a single silicon chip can be used to make thousands of agarose microarrays. The size of the agarose chip can be easily adjusted by making different size of PDMS chips for a wide range of input sample size and concentration. It is also time-saving since it only takes minutes to make an agarose chip for each experiment.

### Cross-species experiments proves the fidelity of Microwell-seq in capturing single-cell transcriptome

The whole workflow of Microwell-seq is shown in Figure 2A. An agrose plate with tens of thousands of microwells is used for trapping thousands individual cells and beads, similar to the Cytoseq platform (Fan et al., 2015). After cells are loaded to the wells, we will inspect the microwell array under microscope and pick out rare cell doublets by capillary tube, which takes about several minutes. Then barcoded magnetic beads are loaded and trapped into each well by size. Every single bead is conjugated with millions of oligonucleotides which share the same cellular barcode (Figure S1). After incubation of beads and cells in a soft flow of lysis buffer, beads with captured mRNA are retrieved by magnet. The procedure from cell loading to the cell lysis can be limited to only 15 minutes, largely preserving the healthiness of the cells. We collect the beads in a 1.5 ml tube where reverse transcription and template switch are performed using Smart-seq2 protocol (Picelli et al., 2013). Amplified cDNA will be fragmented by a customized transposase which carries two identical insertion sequences (see online method). The 3’ ends of transcripts are then enriched in the following library generation PCR and sequenced using Illumina Hiseq platforms (Table S2).

**Figure 2.**
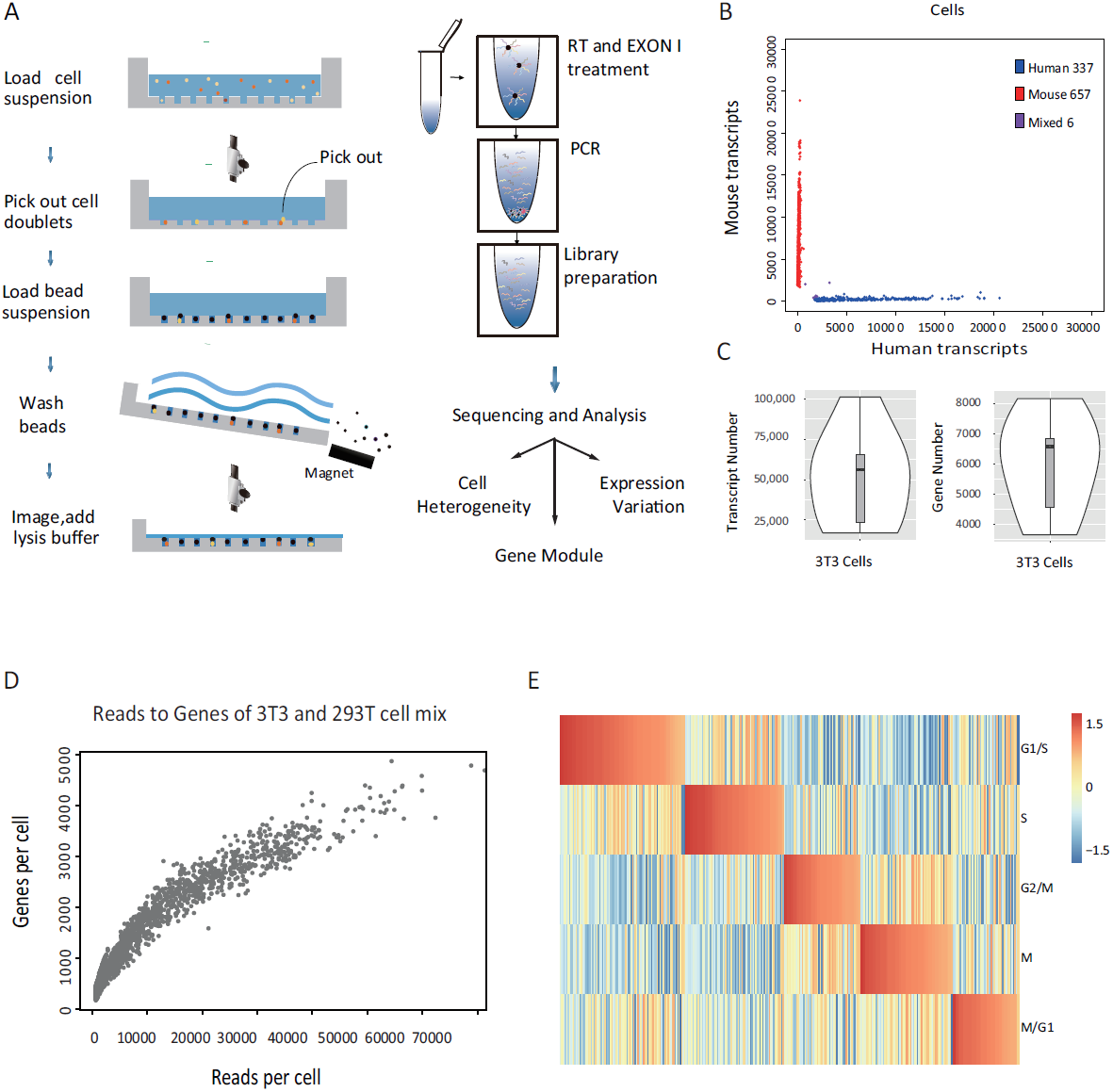
Workflow and Evaluation of Microwell-seq. (A) A schematic of the basic workflow for microwell-seq. (B) Human-mouse mix test by Microwell-seq. Human (293T) and mouse (3T3) cells were mixed at equal concentrations. The scatter plot shows the number of human and mouse transcripts associating to each cellular barcode. Blue dots indicate single cells that were human-specific; red dots indicate single cells that were mouse-specific. 0.6% cell-cell doublets (purple dots) are human-mouse mixed cells. (C) The distribution of gene number and transcripts number in 9 deeply sequenced beads (3T3 cell captured) were shown by violin plots. (D) Detected gene number versus reads number of each individual cell in the species-mixing experiment. (E) Cell cycle state of 337 human (293T) cells measured by Microwell-seq. The score for each phase was calculated using previously reported method (see supplementary data analysis method), and the cell ordered by their phase scores. See also Figure S2 and Table S2.

To evaluate the cross-contamination rate in Microwell-seq method, we performed mixed species experiment with cultured human (293T) and mouse (3T3) cells. Then we identified the ratio of reads mapped to both human and mouse genome in every single cell (Figure 2B). We found that Microwell-seq produced high-fidelity single-cell libraries with no more than 0.6% cell doublets. 8,000 genes and 50,000 transcripts can be detected on average by saturating sequencing (Figure 2C). Low reads versus genes number ratio were observed in large-scale experiments (Figure 2D). Cell cycle score were calculated for each human 293T cell based on previously reported phase-specific genes and methods (Figure 2E) (Macosko et al., 2015; Whitfield et al., 2002). Cells in different cycles are clearly separated based on their cell cycle scores. Cell cycle related gene like *TK1, CDC20, TYMS, UBE2C, CDC20, CCNB1, KPNA2, PCNA* were used to distinguish five phases of cell cycle.

To test the ability of Microwell-seq for profiling hematopoietic cells, we analyzed 4323 thawed single cells of CD34^+^ and CD34^−^ compartments from mobilized human peripheral blood sample. We observed a clear distinction between the two populations with CD34 only expressed in CD34^+^ cells, and various subpopulations were also observed within the two main clusters (Figure S2A). Thawed mPB CD34^+^ cells from batch1 and batch2 were visualized on a t-SNE map. It is worth emphasizing that Microwell-seq work stably even with thawed cells (Figure S2B).

### A transcriptome atlas of 44,914 human HSPCs at single-cell resolution revealed a revised human hematopoiesis model

By harnessing the power of Microwell-seq, we aimed to systematically profile an atlas of adult human stem and progenitor cells with minimal influence from classical flow-based cell type definition system. We therefore collected fresh peripheral blood from two Chinese male donors after GCSF mobilization, and then enriched hematopoietic stem and progenitor cells using CD34^+^ selection kit. FACS analysis showed that 98% of enriched cells were indeed CD34^+^ (Figure 3A). Using Microwell-seq, single cell transcriptomes of 44,914 mPB CD34^+^ cells from 7 independent library generation experiments were successfully profiled. The data is of high quality and is by far the most comprehensive single cell transcriptome atlas of HSPCs.

**Figure 3.**
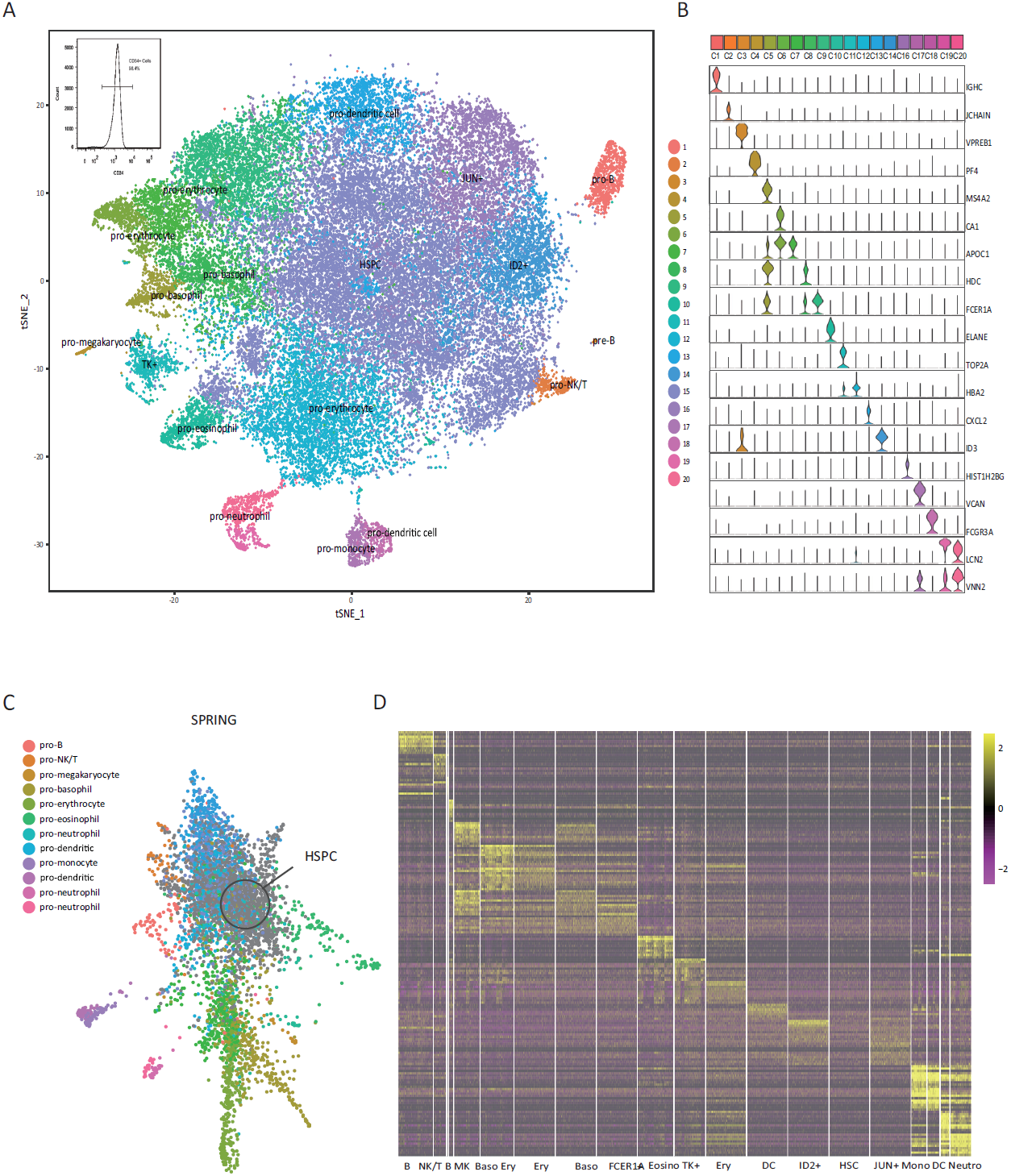
Analysis of adult human hematopoietic stem and progenitor cells by Microwell-seq. (A) t-Distributed Stochastic Neighbor Embedding (t-SNE) analysis of 44,914 mPB CD34^+^ cells. 20 subpopulations were defined and labeled in the t-SNE map. (B) Expression pattern of candidate genes among 20 inferred cell populations. Violin plots were used to reflect the distribution of gene expression level within each cluster. (C) 4289 cells from batch 4 were presented on the two-dimension space reduced by SPRING. SRPING depicts the dynamic trajectories of progenitor cells (color labeled) that differentiate from HSPCs (labeled as grey). (D) Differentially expressed genes across 20 cell populations. Each cluster was downsampled to no more than 1000 cells to prevent dominating the plot. Rows correspond to individual genes found to be selectively upregulated in individual clusters; columns are individual cells, ordered by cluster (1–20). See also Figure S2, S3 and Table S3.

Through computational analysis (Satija et al., 2015), we identified 20 subpopulations (Figure 3A). Each subpopulation showed a highly specific gene expression pattern (Figure 3B). We found that, within the same donor, batch effects from different library generation experiments were minimal, as all 20 clusters were reproducibly seen from all 7 batches of experiments, with slight proportion difference between the two assayed donors (Figure S2C and S2D). From down sampling experiments, we found that sample size should reach ten thousand single cells to find major defined subpopulations (Figure S3). 4289 cells were presented on the two-dimension space reduced by SPRING (Weinreb et al., 2017), depicting the dynamic trajectories of progenitor cells that differentiate from HSPCs (Figure 3C).

Heatmap showed the hierarchical clustering of the single cell data sets with clear differential gene expression modules and cell type clusters (Figure 3D and Table S3). Interestingly, many of the defined human myeloid clusters such as the *CEBPE*^+^ cluster, *CD14*^+^ cluster and *CD41*^+^ cluster, strongly resemble published mouse myeloid clusters by MARS-seq analysis (Jaitin et al., 2014; Paul et al., 2015), suggesting that these clusters are conserved among species. The related markers of these clusters are also highlighted in another recent single cell analysis work on human hematopoietic progenitors using a MARS-seq like platform (Velten et al., 2017), indicating consistency of the finding from various technologies.

We found progenitors primed with unique lineages like megakaryocyte, neutrophil and B cell progenitor can project directly from HSPCs. C1, C4, C5, C6, C10, C19, C17 and C18 are distinguished by the differential expression of *MS4A1, PF4, LMO4, TFRC, ECP, CEBPE, CSF1R* and *CD74*, which are associated with known signatures of B cell, megakaryocyte, basophil, erythrocyte, eosinophil, neutrophil, monocyte and DC progenitors, respectively.

We can see a developmental continuum of B cell differentiation in clusters C3-to-C1. The significant expression of *VPREB1 and VPREB3* suggested C3 as the B precursor progenitor. *CD79A* and *CD79B* were high expressed in C1, indicating its B cell progenitor identities. B cell related genes, including *MS4A1, IGHD* and *CD19*, were also found in C1 (Novershtern et al., 2011). Highly expressed marker *NKG7, DNTT and ACY3*, together with TFs *MLXIP*, defined C2 as natural killer and T cell (NK/T) progenitor (Novershtern et al., 2011; Zheng et al., 2017). Cluster 4 (C4) was identified by high expression of *ITGA2B (CD41), CD9, PF4, PPBP* and *PLEK*, which suggests a megakaryocyte fate (Notta et al., 2016; Paul et al., 2015). C5/C8 was defined as basophil progenitor by the expression of transcription factors (TFs) *GATA2* and *LMO4*, along with basophilic gene like *MS4A2, TPSB2* and *RNF130* (Paul et al., 2015). Markers of erythrocyte progenitors, including *GATA1, KLF1, MYC, TFRC (CD71), HBB* and *HBD*, were strongly expressed in C6 and C7, which delineated them as CD34^+^ erythrocyte progenitors (Novershtern et al., 2011; Paul et al., 2015). *GATA1* and *KLF1* play well-established roles in erythroid differentiation (Novershtern et al., 2011). In addition, C12 showed high *HBB* and *SOX4* expression, indicating its erythroid identity (Paul et al., 2015). These results provided powerful support for the existence of erythroid progenitors in the CD34^+^ compartment, consistent with the study of Notta et al. (Notta et al., 2016).C10 and C19/C20 were defined as eosinophil and neutrophil progenitors: C10 was marked by expression of RNase2 *(EDN)* and RNase3 *(ECP)* (Rosenberg et al., 2013) , which suggested an eosinophil fate. They also co-expressed granulocyte genes like *CST7* (Paul et al., 2015). C19 showed early neutrophil characteristics, with extremely high level of *CEBPE*, along with *CSF3R (CD114), CD1*77, *S100A9, S100A8* and *S100P* (Novershtern et al., 2011; Paul et al., 2015). C20 showed similar pattern to C19 but with lower gene expression level. C17 exhibited strong levels of known monocyte markers including *LYZ, CD14, CSF1R, MS4A7*, and monocyte TFs like *FCN1* and *MNDA* (Novershtern et al., 2011; Paul et al., 2015). C18 was defined as dendritic cell progenitor by expression of *CD74*, MHCII-related genes and the TFs *STAT1, ID2* and IRF8.C13 was suspected to be epidermal/Langerhans dendritic cell progenitor by the expression of *CXCL2*, which was closely related to dendritic cell maturation (Lee et al., 2009). *ID2*, enriched in C14, was very recently regarded as an essential marker of dendritic cell (Paul et al., 2015). C11 expressed high levels of cell cycle related gene like *TK1* and *PCNA*. C16 uniquely expressed *JUN*, which should be early T cell characteristic (Novershtern et al., 2011).

Finally, we found that C15 was different from all the other clusters by lack of any lineage-defining TFs or markers (e.g. *MNDA /KLF1/GATA1* or *ELAN /CD62L /MPO)*. These cells did express HSPC markers such as *CD34, SPINK2* and *SOX4*, so we define C15 as the undifferentiated multipotent hematopoietic stem and progenitor type in agreement with Paul et al. study (Paul et al., 2015).

Based on single cell transcriptome profiling, our analysis suggests that the classically defined oligopotent progenitor types such as common myeloid progenitor (CMP), common lymphoid progenitor (CLP), megakaryocytic/erythroid progenitor (MEP), granulocyte and monocyte progenitor (GMP), common DC precursor (CDP), macrophage/DC progenitor (MDP) and multilymphoid progenitor (MLP) cannot be recognized in the main clusters of adult human hematopoietic stem and progenitor cells (Doulatov et al., 2010; Geissmann et al., 2010; Novershtern et al., 2011; Onai et al., 2007). On the other hand, we found that the mPB CD34^+^ cells are dominated by progenitors with expression of lineage specific modules. Interestingly, even in the multipotent cluster C15, we still observe significant heterogeneity, suggesting a more complexity within hematopoietic stem cells (Figure S4). In the adult human hematopoietic hierarchy, unipotent progenitors appear to be directly downstream of multipotent progenitors. The oligopotent intermediate state cannot be found from the transcriptome-based single cell analysis.

### Extraction of subcluster structure within the defined progenitor types

Using the comprehensive data set, we then aim to further dissect the subcluster structure within the defined progenitor types. All defined clusters show further heterogeneity, suggesting the complexity of the differentiation pathway (Figure 4A). We found the cluster 1 can be further grouped into three subsets, each with specific gene expression pattern. Subset 1 of C1 (C1S1) is enriched for *CD1C, IGLC3* and *CLECL1*, C1S2 highly expresses *TCL1A, TXNIP* and *IGDH* gene cluster, and C1S3 expresses *TXN, IGLL* and *NUCB2*. There were also three subsets with distinct expression pattern in cluster 2: C2S1 with gene *NLY* and *PLD4*, C2S2 with gene *TBC1D7* and *AKAP13*, C2S3 with gene *SPON1* and *RGS1*. Cluster 4 can be divided into two groups, one highly expresses the *ERH, NCL* gene cluster, and the other expresses the *ACRBP, MMD, TUBB1* gene cluster. Meanwhile, there were three subsets with distinct gene expression in cluster 10: C10S1 high expressed gene *LY86* and *VCAN*, C10S2 enriched for gene *CALR* and *P4HB*, and C10S3 accompanied with gene *CD52* and *TRBC2*. In addition, we found that cluster 19 can be further grouped into 4 subsets: C19S1 highly expresses gene *PI3* and *VNN2*, C19S2 enriched for gene *EGFL7* and *CDK6*, C19S3 accompanied with gene *CAMP andTCN1*, and C19S4 expresses genes such as *DEFA4, AZU1*, and *SLP1*.

**Figure 4.**
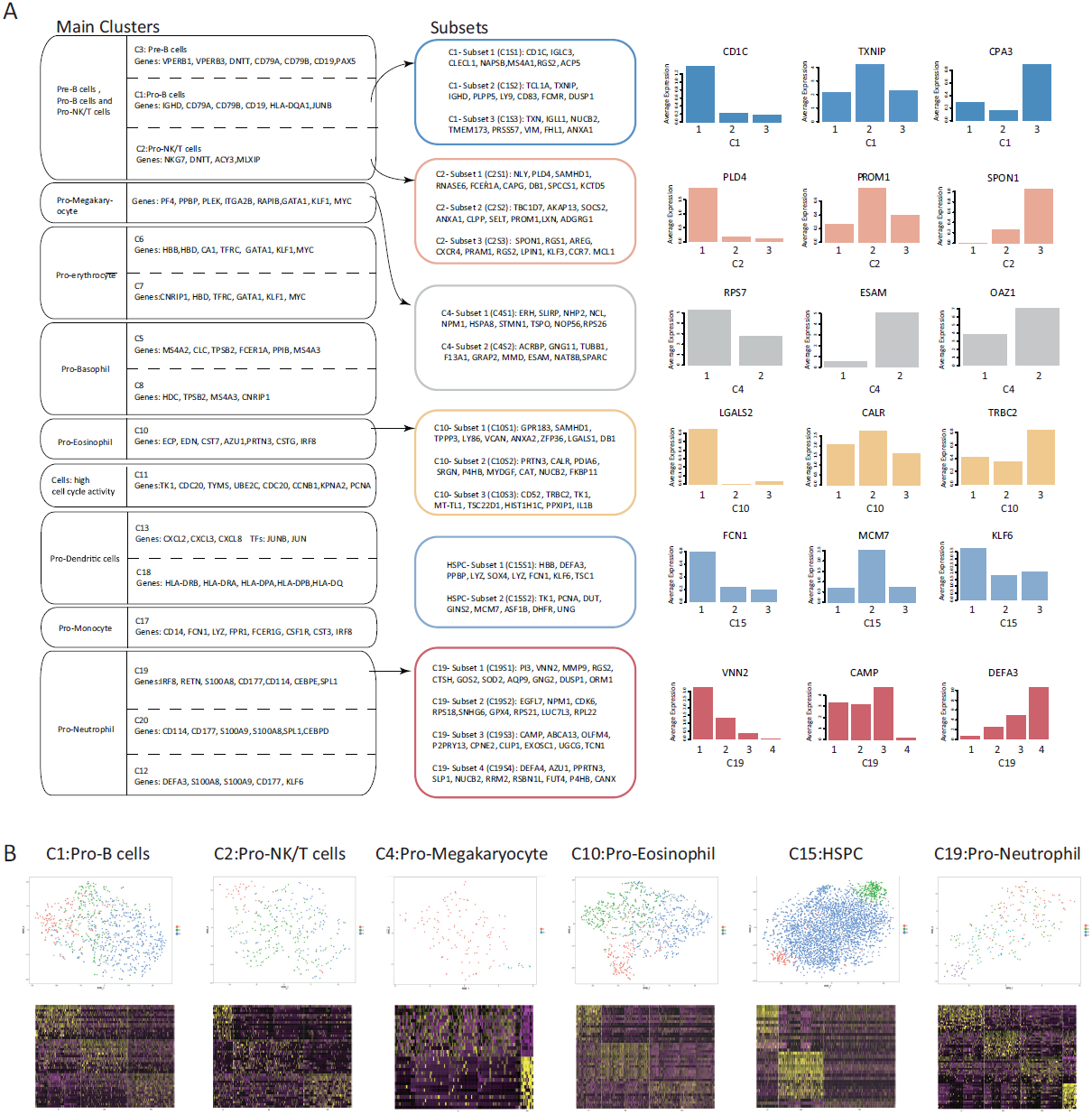
Extraction of subsets from defined progenitors. (A) Gene expression pattern of main progenitor types as well as their subsets. All subsets have unbiased cluster gene expression but unique and specific subset gene expression. The bar charts show the average expression level of selected genes in each subsets. (B) tSNE plots and heatmaps showing expression pattern of different main progenitor types and their subsets. Subcluster structure of cluster 1, 2, 4, 10, 15 and 19 were chosen as represents. See also Figure S4.

It worth mentioned that we found three subgroups even in the defined HSPC population. C15S1 appears to correspond to an early erythroid and megakaryocytic progenitors with higher expression of *HBB* and *PPBP*. C15S2 is enriched for some unknown gene like *DUT, GINS2* and cell cycle related gene including *TK1* and *PCNA*. Finally, C15S3 is not characterized by any lineage specific marker suggesting a more primitive HSPC state. The subcluster structure of different populations appears to correspond to different differentiation stages rather than contaminations and technical noise. Based on the clear cell clusters and gene expression modules, we infer that the human hematopoietic lineage differentiation takes concrete but unidirectional “steps” to move from multipotent stage to mature cell types (Figure 4B).

### Modeling lineage differentiation pathway in the human hematopoietic system

In order to obtain the reference single cell transcriptomes for major differentiated cell types in the adult human hematopoietic hierarchy we further profiled 4900 CD34^−^ cells from GCSF mobilized peripheral blood sample. We defined eight main clusters within our CD34^−^ dataset. After downsampled CD34^+^ dataset to the same scale as CD34^−^ dataset, we visualize it on a 2-dimension tSNE map (Figure 5A). mPB CD34^+^ cells and mPB CD34^−^ cells were separated. The main force driving them apart results from distinct transcriptional expression. mPB CD34^+^ cells have higher level of CD34 expression, further convincing its identity as progenitor cells (Figure S5A). All cells were divided into 28 clusters. P1-P20 is CD34 positive cells that above mentioned C1-C20, including pro-eosinophil, pro-neutrophil, pro-natural killer cells, pro- B cell, pro-megakaryocyte, pro-erythrocyte, pro-NK/T, pro-basophil, pro-dendritic cells and pro-monocytes (Figure 5B). Meanwhile, mature cell (CD34^−^) cluster N1-N8 correspond to eosinophil, neutrophil, NK/T cells, B cell, dendritic cells and monocytes (Figure 5C). The differential markers within the 8 CD34^−^ clusters were also found in CD34^+^ clusters, linking progenitors and mature cells from these two datasets.

**Figure 5.**
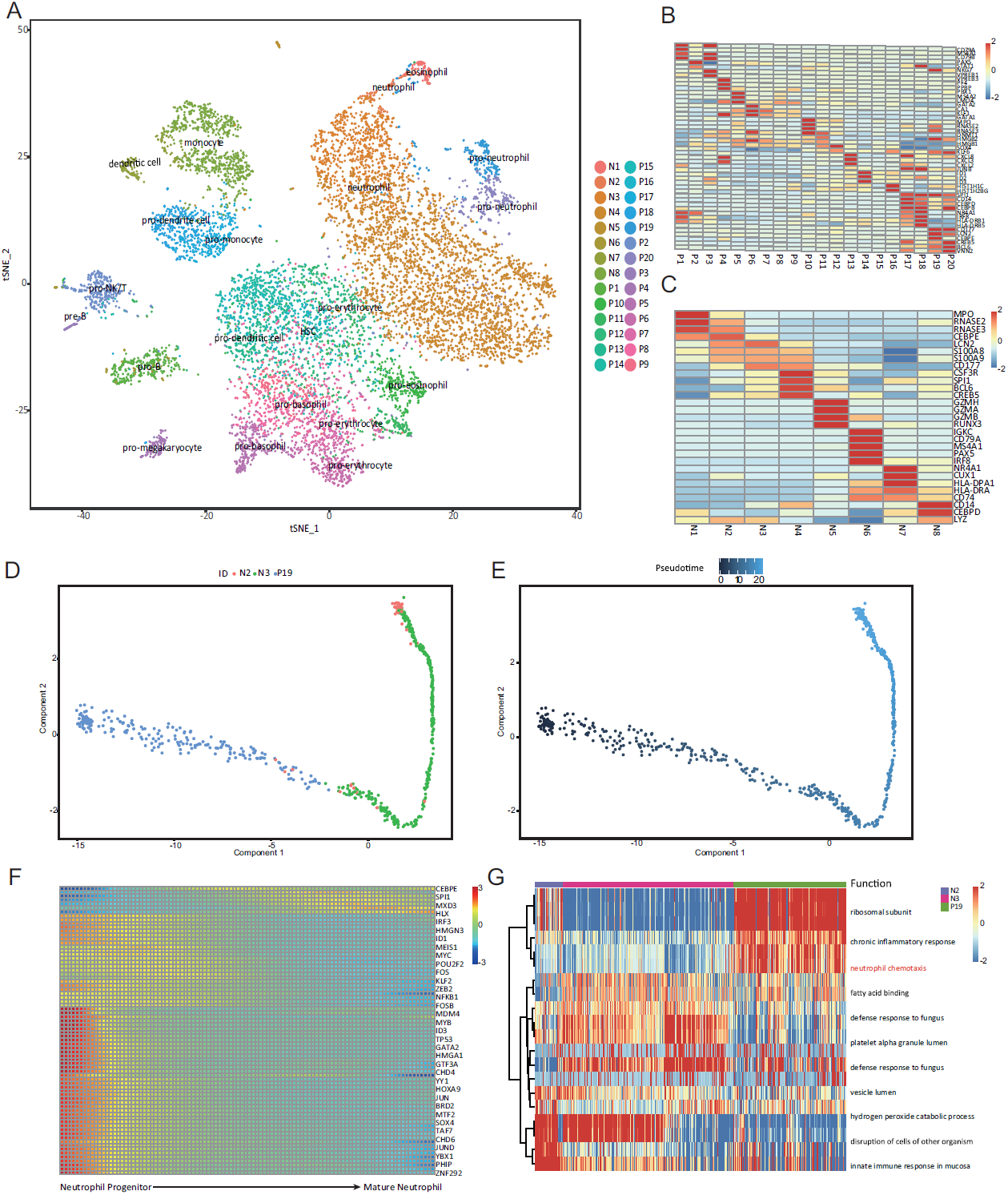
Integration of data from CD34^−^ and CD34^+^ hematopoietic cells. (A) 5512 mPB CD34^+^ cells and 4973 mPB CD34^−^ cells were visualized on the t-SNE map. Cells were labeled by predefined cell type identity. N1-N8 represent eight main clusters in mPB CD34^−^ cells while P1-P20 represent twenty main clusters in mPB CD34^+^ cells. (B) and (C) heatmaps show mean expression per cluster within CD34^−^ and CD34^+^ two datasets for lineage specific TFs and markers. (D) and (E) pre-clustered cells that are associated with neutrophil from CD34^−^ and CD34^+^ combined dataset are plotted on the 2-dimension space reduced by Monocle. Cells are labeled as its identity and pseudotime, respectively. (F) Differentially expressed TFs along the pseudotime line. Cells are ordered according to its pseudotime (from left to right). Not all gene names are listed in figure. (G) Differentially expressed gene sets named by biological processes. Gene sets and cells are clustered using Euclidean distance. Not all biological processes’ names are shown in figure. See also Figure S5.

N1, highly expressing *EDN* and *ECP* and granulocyte marker *MPO*, is mature eosinophil, corresponding to the P10. N2-N4 defined by neutrophil-specific makers like *CEBPE, CD177* and *CSF3R (CD114)*, is relevant to P19-20. Markers of NK cell, including *GZMH, GZMA* and *GZMB*, were strongly expressed in N5, which delineated them as mature NK cells. N5 also highly expressed T cell markers such as CD3, CD8, CXCR4 and CD69 (Bagger et al., 2016). So we speculated that N5 were NK/T cell, a heterogeneous group of T cells that share properties of both T cell and NK cell (Godfrey et al., 2004). N5 is related to P2, The NK/T progenitor. B cell markers gene like *CD79A, IGKC* and *MS4A1* are enriched in N6, linking them to P1, the B cell progenitor. N7 was marked by expression of *CD74* and MHCII related gene, which suggested a dendritic fate, corresponding to P18. N9 is closely related to P17, showing monocyte characteristics, with high expression of *CD14*, along with *LYZ* and *CEBPD* (Novershtern et al., 2011; Paul et al., 2015). Correlation heatmap also indicated a partial similarity between CD34^+^ clusters and CD34^−^ clusters (Figure S5B). Selecting neutrophil as a candidate, we established pseudotime from neutrophil progenitor cells to mature neutrophil cells (Figure 5D and 5E) and constructed transcriptional priming pathways in neutrophil cell (Figure 5F and 5G). Transcription factors such as *CEBPE* and *SPL1* gradually increased from progenitor cells to mature cells, while transcription factors such as *PHIP* gradually decreased.

### Validation of the revised hematopoietic differentiation model

To test the stability of the revised hematopoietic differentiation model, we transplanted 1×10^5^ mPB CD34^+^ cells into sublethally irradiated NCG mouse (NOD-Prkdc^em26Cd52^Il2^rgem26Cd22^/Nju, Nanjing Biomedical Research Institute of Nanjing University) via injection into the femur cavity. Two months after transplantation, human CD34^+^hCD45^+^ cells in mice femur and tibia were sorted by flow cytometry using anti-human-CD45-PE and anti-human-CD41a-FITC antibodies. We found that the multipotent to unipotent hierarchy in the human hematopoietic system is largely maintained in the xenograft experiment. Clustering Heatmap shows the distinct gene expression pattern of pro-neutrophil, pro-megakaryocyte, pro-T, pro-B and pro-Ery modules (Figure 6A). Examples of xenograft-stable progenitors were labeled in t-SNE map (Figure 6B).

**Figure 6.**
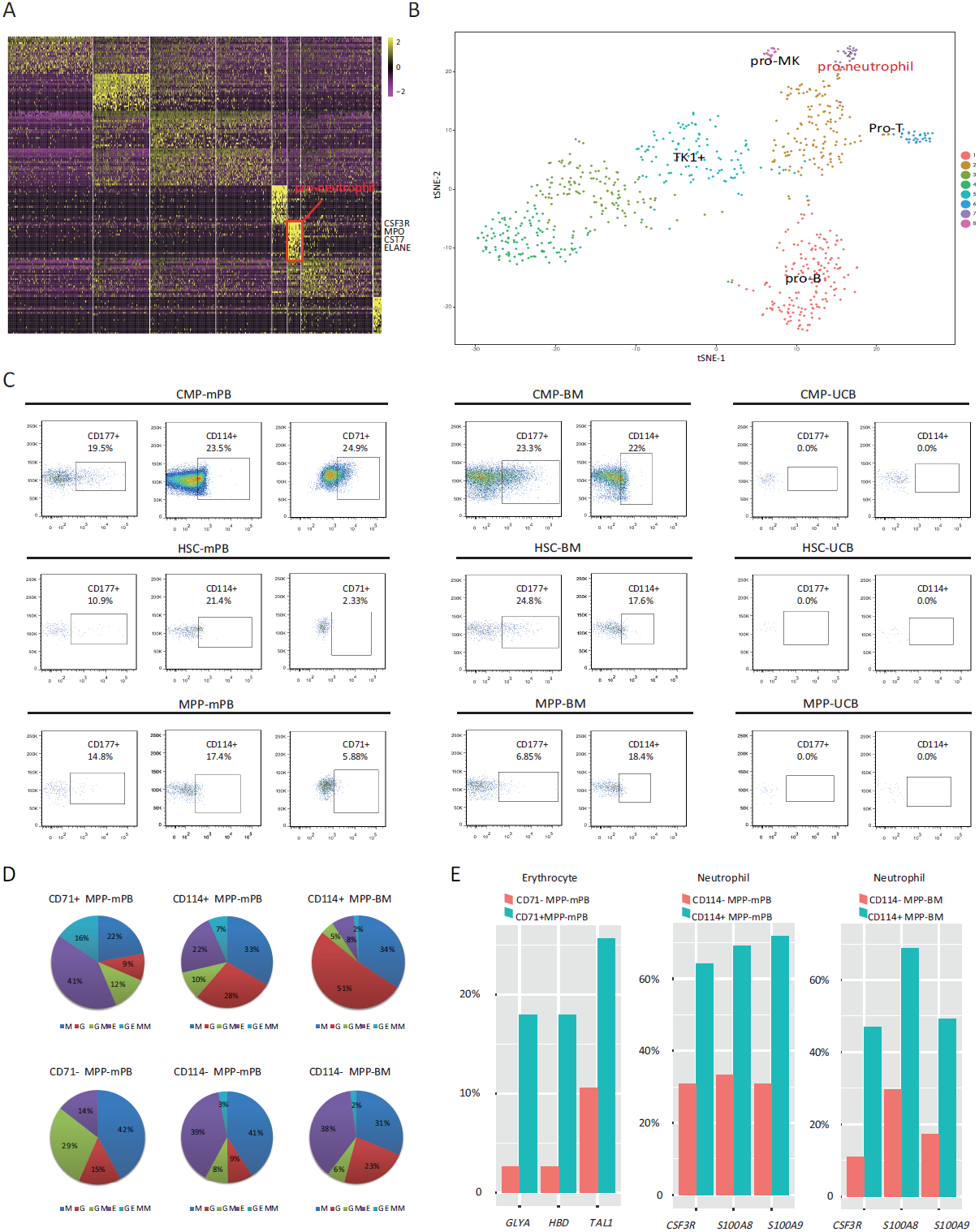
Functional validation of the revised adult human hematopoietic hierarchy. (A) A clustering Heatmap showing the gene expression pattern of CD34^+^hCD45^+^ cells sorted from mouse bone marrow two weeks after transplantion. (B) Lineage primed progenitors like pro-neutrophil, pro-megakaryocyte pro-T and pro-B in CD34^+^hCD45^+^ cells were visualized on the t-SNE map.(C) The gating scheme of defined Neutrophil progenitors (CD114^+^/CD177^+^) from CMP(CD34^+^CD38^+^CD45RA^−^), HSC(CD34^+^CD38^−^Thy1^+^CD45RA^−^CD49f^+^) and MPP (CD34^+^CD38^−^Thy1^−^CD45RA^−^CD49f^−^). Erythrocyte progenitors (CD71^+^) from CMP(CD34^+^CD38^+^CD45RA^−^) and MPP(CD34^+^CD38^−^Thy1^−^CD45RA^−^CD49f^−^). CMP, HSC and MPP were sorted from peripheral blood cells, bone marrow cells and umbilical cord blood cells by FACS. (D) The percentage of different colony types in the colony forming assay from CD71^+^ and CD71^−^ MPP-mPB, CD114^+^ and CD114^−^ MPP (mPB and BM). The results showed the average percentage from 3 independent experiments. CD71^+^ MPP-mPB were enriched for CFU-E making cells when compared with CD71^−^ MPP-mPB. CD114^+^ MPP (mPB and BM) were enriched for CFU-G making cells when compared with CD114^−^ MPPs. (E) Single-cell qPCR was used to detect the expression of erythrocytes and neutrophils markers in the cells from colony forming assay. 39 cells from CD71^+^ MPP-mPB clones, 38 cells from CD71^−^ MPP-mPB clones, 39 cells from CD114^+^ MPP-mPB clones and 39 cells from CD114^−^ MPP-mPB clones were picked out for Biomark single cell qPCR. 64 single cells for CD114^+^ bone marrow MPP derived clones and 128 single cells for CD114^−^ bone marrowMPP derived clones were randomly picked out for Realtime single cell qPCR. *HBD, TAL1* and *GLYA* were selected as markers for erythrocytes. *S100A8, S100A9* and *CSF3R* were selected as markers for neutrophils. The percentages of differentiated cells expressing erythroid or neutrophil markers were shown in the barchart. See also Figure S6 and Table S4.

To further validate the defined lineage primed progenitors and their early differentiation pattern, we searched for cell surface markers and available antibodies that can best characterize different clusters (Supplementary Material). Considering both cluster specificity and antibody availability, we used CD41, TFRC (CD71), CSF3R (CD114), CSF1R (CD115) and MHCII to mark early megakaryocytic, erythroid, neutrophil, monocytic and dendritic progenitors respectively. With specific antibodies against these markers, we could separate at least 5 kinds of lineage biased progenitors from human MPP (CD34^+^CD38^−^Thy1^−^CD45RA^−^CD49f^−^) cells, HSC (D34^+^CD38^−^Thy1^+^CD45RA’CD49f^+^) and CMP (CD34^+^CD38^+^CD45RA^−^) cells respectively (Figure 6C and Figure S6).

The gating scheme defined neutrophil progenitors (CD114^+^/CD177^+^) and erythrocyte progenitors (CD71^+^) from CMP, HSC and MPP sorted from peripheral blood cells, bone marrow cells and umbilical cord blood cells. In the mobilized peripheral blood (mPB) derived CMP, there were 19.5% CD177^+^ neutrophil progenitors, 23.5% CD114^+^ neutrophil progenitors and 24.9% CD71^+^ erythrocyte progenitors. Meanwhile bone marrow (BM) derived CMP presented a similar proportion, CD177^+^ neutrophil progenitors and CD114^+^ neutrophil progenitors were 23.2% and 22% respectively. Then we search progenitors at HSC level, the peripheral blood-derived group have 10.9% CD177^+^ neutrophil progenitors, 21.4% CD114^+^ neutrophil progenitors and 2.33% CD71^+^ erythrocyte progenitors, while bone marrow-derived group have 24.8% CD177^+^ neutrophil progenitors, 17.6% CD114^+^ neutrophil progenitors. Finally, we found progenitors even exist in MPP cells. In the peripheral blood-derived MPP, there were 14.8% CD177^+^ neutrophil progenitors, 17.4% CD114^+^ neutrophil progenitors and 5.88% CD71^+^ erythrocyte progenitors. In addition, bone marrow-derived MPP showed a similar proportion, CD177^+^neutrophil progenitors and CD114^+^ neutrophil progenitors were 6.85% and 18.4% respectively. It is worth mention that neutrophil progenitors (CD114^+^/CD177^+^) were absent in CMP, HSC and MPP derived from umbilical cord blood (UCB).

Single cell clonal forming assay showed that CD71^+^ selection remarkedly enriched CFU-E generating cells. 41% of clones generated by CD71^+^ MPP-mPB were CFU-E, while only 11% of clones were found to be CFU-E for CD71^−^ MPP-mPB. In addition, CD114^+^ selection enriched CFU-G generating cells. The percentage of CFU-G generating cells in CD114^+^ MPP-mPB is three times as large as that in CD114^−^ MPP-mPB. Similarly, 51% of clones produced by CD114^+^ MPP-BM were CFU-G, while only 23% of clones were found to be CFU-G for CD114^−^ MPP-BM (Figure 6D). We used single cell qPCR assay to analyze gene expression of single cells randomly picked up from those clones (Table S4). The percentages of cells highly expressing *TAL1, HBD* and *GLYA* were substantially higher within the clones formed by CD71^+^ MPP-mPB as compared with CD71^−^ MPP-mPB. The percentages of cells highly expressing *CSF3R (CD114), S100A8* and *S100A9* were higher within clones from the CD114^+^ MPP (mPB and BM) than CD114^−^ MPP (mPB and BM) (Figure 6E). These results suggest both CD71 and CD114 are able to enrich lineage biased progenitors from MPP and lineage differentiation begins in early MPP stage. Thus, we provide functional evidence that the lineage primed progenitors in the adult human hematopoietic hierarchy are significantly biased for their lineage potential. We also identify a novel neutrophil differentiation pathway that is independent of classical myeloid progenitors.

## Discussion

The method used in our study holds many advantages over other related technologies (Fan et al., 2015; Gierahn et al., 2017; Klein et al., 2015; Macosko et al., 2015; Yuan and Sims, 2016). Microwell-seq demonstrates superiority in cost and convenience. A silicon wafer containing ~100,000 microwells cost $ 700, can be used to make hundreds of polydimethylsiloxane(PDMS) micropillar array, and a PDMS array can be used repeatedly to produce hundreds of agarose-made microwell array. The cost per experiment is no more than the cost of agarose, which is negligible. The magnetic propertiy of barcoded beads makes the collection efficient. The extra beads outside the microwells can be retrieved by a magnet. In addition, cell doublets can be picked out by micromanipulation, which ensures Microwell-seq to produce high-fidelity single-cell libraries with no more than 0.6% cell doublets rate, much lower than other platforms. By reducing the size of the PDMS chip, our agrose micowells works perfectly with limited number of precious cells. Finally, because the experimental devices requires nothing more than an ordinary microscope, it is very easy to scale up the experiment by handling multiple microwells at the same time. Thus, Microwell-seq is portable, efficient, faithful, and cheap high-throughput scRNA-seq platform.

Using Microwell-seq, we presented a comprehensive revised model of complex hematopoietic hierarchy. Through the transcriptomic analysis of around 50,000 single adult human hematopoietic cells, we found 28 subpopulations including pro-eosinophil, pro-neutrophil, pro-natural killer cells, pro- B cell, pro-megakaryocyte, pro-erythrocyte, pro-NK/T, pro-basophil, pro-dendritic cells and pro-monocytes and related mature cell types like eosinophil, neutrophil, NK/T cells, B cell, dendritic cells and monocytes. Meanwhile, we demonstrated the gene clusters that distinguish each progenitor. Compared those with mature cells, we confirmed the lineage primed modules that we have defined with the single cell transcriptome analysis are not a result of transcriptional or technical noise. The transcriptional priming occurring at the early progenitor stage will lead to a biased lineage decision, challenging classical model of hematopoiesis. Our results strongly support a revised adult human hematopoietic differentiation pathway independent of oligopotent progenitors. Neutrophil progenitor was chosen as a candidate to dissect the transcriptional priming factors and pathway transition from priming state to mature state. Our study also resolves the maturation trajectory from neutrophil progenitor to mature neutrophil lineages.

Our results provide strong supports for recently published single cell analysis work with mouse and human hematopoietic systems (Notta et al., 2016; Paul et al., 2015; Velten et al., 2017). We observe an HSPC population that strongly resembles the “CLOUD-HSPC” proposed by Velten et al. (Velten et al., 2017). We observed all the described early myeloid differentiation pathways proposed by Notta et al., Paul et al. and Velten et al. studies. However, our work differs from these studies in the following aspects. First, we did not use FACS to enrich specific progenitor types and ensured an unbiased and comprehensive analysis for all of the adult human hematopoietic stem and progenitor populations. Secondly, the throughput of our analysis is higher than all of the published work by more than an order of magnitude; the processed cell number of the microwell-seq experiment is now close to normal FACS analysis. Thirdly, the comprehensiveness of our single cell atlas data resource for the human hematopoietic stem and progenitor cells allowed us to find many unknown clusters that are interesting for future studies. Finally, we validated an early neutrophil differentiation pathway that is previously uncharacterized.

We found that *CD114* is found to be highly heterogeneous in adult human CD34 ^+^ cells. Moreover, CD114 antibody enriches neutrophil progenitors based on our single cell clonal forming assay and single cell qPCR assays¨ Interestingly CD114^+^ pro-neutrophils can be found in both adult marrow and mobilized peripheral blood, but are absent in umbilical cord blood. Umbilical cord blood cells are becoming a prominent source of cells for HSC-based therapeutics due to the shortage of adult bone marrow HSCs and mobilized peripheral blood. However, it is well known that the time to neutrophil recovery, a major indicator of posttransplant mortality, is longer for cord blood than adult marrow (Brunstein and Wagner, 2006). This phenomenon affects the efficacy of transplantation to some degree. The existence of neutrophil progenitor in adult HSPCs may be responsible for this phenomenon, which is worth for future investigation.

In conclusion, Microwell-seq opens a way for all laboratories around the world to harness the power of single-cell RNA-seq technology. We believe this easy to use system will gain its popularity and benefit the biomedical research community in the near future. In addition, the revised hematopoietic hierarchy model will provide guidance for the clinical implementation of HSCs transplantation and scientific research in hematology.

## Supplemental Information

Supplemental Information includes 6 figures, 4 tables and a supplemental data analysis method.

Data Access

Digital Expression Matrixes of all samples are accessible on the following link: https://figshare.com/s/4cb39e67615c8154717b

## ACKNOWLEDGMENTS

We thank L. Shen, S. Ying, B. Liu, F. Tang, S. Quake, G. Fan, X. Li, X. Yu, L. Sun for help on experiments. We thank G-BIO, Novogene, Annoroad and Geneseeq for deep sequencing experiments. We thank Vazyme for supplying customized enzymes for the study. This work was supported by funding from Fundamental Research Funds for the Central Universities (G.G.) and 1000 Youth Talent Plan (G.G.). G.G. is a professor from Zhejiang University.

## CONTRIBUTIONS

The project was designed by G.G.; S.L. and W.H. contributed to the development of Microwell-seq; Experiments were carried out by S.L., M.J., H.C., F.Y., R.W., X.J., D.H., Y.Q., J.M. and X.H.; Sequencing data was analyzed by Y.X. and W.H.; The paper was written by S.L., W.H., Y.X. and G.G. This work was supported by Zhejiang University Stem Cell Institute.

## COMPETING FINACIAL INTERESTS

G.G. has submitted a patent application related to the Microwell-seq method reported in this paper.

## Online method

### 3T3 and 293T preparation

293T and 3T3 cells were grown in DMEM (Corning, cat #10-013-CVR) with 10% FBS (Life Technologies, cat #10099141) and 1% penicillin-streptomycin (Life Technologies, cat #15140122). Cells were washed twice by DPBS (Corning, cat #21-031-CVR), then treated with 0.25% trypsin (Invitrogen, cat #12604013) and quenched with growth medium. Cells were diluted by 1x DPBS (Corning, cat #21-031-CVR) with 2mM EDTA (Life Technologies, cat #15575020) and passed through a 40-micron cell strainer (Biologix, cat #151040). Cells were counted by hemocytometer and diluted to ~100,000/mL in 1x DPBS with 2mM EDTA.

### Human CD34^+^ mobilized peripheral blood collection and process

Human mobilized peripheral blood samples were collected from donors at The First Affiliated Hospital of Zhejiang University School of Medicine (Zhejiang, China). The donors have signed the informed consent form. All procedures are approved by the Ethical Committee on Medical Research at School of Medicine. mPB CD34^+^ cells were selected by human CD34 selection kit (StemCell Technologies Cat #18056), following the protocol provided by the manufacturer. After spin at 300 × g for 5min, the supernatant was removed and mPB CD34^+^ cells were suspended in 1× DPBS ^+^ 2mM EDTA. For Microwell-seq, cells were diluted to ~ 100,000/mL.

### Immunodeficient mouse transplantation

mPB CD34^+^ cells were transplanted into sublethally irradiated NCG (NOD-Prkdc^em26Cd52^Il2rg^em26Cd22^/Nju, Nanjing Biomedical Research Institute of Nanjing University) mice via an injection into the femur cavity (approximately, 1×10^5^ cells per mouse). Two months after transplantation, CD34^+^hCD45^+^ cells were sorted from mice bone marrow cells collected from the femur and tibia. NCG mice were fed in Laboratory Animal Center of Zhejiang University and all procedures are approved by the Ethical Committee on Laboratory Animal Center of Zhejiang University.

## Microwell-seq procedure

### Cell collection and lysis

Cell concentration should be carefully controlled in Microwell-seq. Both cell and bead concentrations were estimated by hemocytometer. The proper cell concentration is ~ 100,000 cells/mL(10% wells will be occupied by single cells). The proper beads concentration is ~ 1,000,000/mL(nearly every well will be occupied by single beads). Evenly distributed cell suspension were pipetted onto microwell array and extra cells were washed away. To eliminate cell doublets, the plate was inspected under a microscope. Cell doublets were removed by a capillary tube. Bead suspension was then loaded on the microwell plate which was placed on a magnet. Excess beads were washed away slowly. Cold lysis buffer (0.1 M Tris-HCl pH 7.5, 0.5 M LiCl, 1% LiSDS, 10 mM EDTA, and 5 mM dithiothreitol) was pipetted over the surface of the plate, and removed away after 10 min incubation. Then beads were collected and transferred to an RNase-free tube, washed twice with 1 mL 6X SSC, and once with 1ml of 1× RT buffer. Finally, beads were evenly distributed into five 1.5mL tubes, and every tube contains ~ 10,000 beads.

### Reverse transcription

This step followed the instructions from Smart-seq2 protocol. 1 μL dNTP mix (10mM) was added to every tube and incubate in room temperature for 1 min. Then 9 μL RT mix was added. The RT mix contained 100U SuperScript II reverse transcriptase, 1x Superscript II first-strand buffer (Takara, cat #2690A), 10 U Rnase Inhibitor (Sangon, cat #B600008), 1M Betaine (Sigma, cat #B0300-1VL) and 6mM MgCl_2_ (Ambion, cat #AM9539G). The beads were incubated at 42°Cfor 90 minutes (shake beads every 15 min), followed by 72°C for 15 minutes to inactivate reverse transcriptase.

### Exonuclease I treatment

After washing by 1X exonuclease I buffer, beads were suspended in 200 μL exonuclease I mix (containing 1x exonuclease I buffer and 1 U/μL exonuclease I (NEB, cat #B0293S)), and incubated at 37°C for 45minutes to remove oligonucleotides which did not capture mRNA. Beads were pooled and washed once with 1 mL TE-SDS (1x TE + 0.5% Sodium Dodecyl Sulfate), once with 1 mL TE-TW (10 mM Tris pH 8.0, 1mM EDTA, 0.01% Tween) and once with 1 mL ultrapure distilled water (Life Technologies, cat #10977015).

### cDNA amplification

Beads were then distributed to 5 tubes. Every tube was added with 25 μL PCR mix, which includes 1× HiFi HotStart Readymix (Kapa Biosystems, cat #KK2602) and 0.4uM TSO_PCR primer (see Supplementary Material). The PCR program was as follows: 98°Cfor 3min, 12 cycles of (98°C20sec, 67°C 15sec, 72°C6min), 72°C5min and 4°Chold. After pooling all PCR products, AMPure XP beads (Beckman Coulter, cat #A63881) were used to purify the cDNA library.

### Transposase fragmentation and selective PCR

Purified cDNA library was fragmented by a customized transposase which carries two identical insertion sequence. The customized transposase was from TruePrep DNA Library Prep Kit V2 for Illumina (Vazyme, cat #TD513). Fragmentation reaction was done following the instructions of manufacturer. We replace the index 2 primers (N5××) in the kit with our P5 primer (see supplementary Material) to specially amplify fragments which contain the 3’ end of transcript. Other fragments will form self-loops, which impede their binding to PCR primers. PCR program is as follows: 72°C3min, 98°C30sec, 14 cycles of (98°C15sec, 60°C30sec, 72°C3min) 72°C5min and 4°Chold. To eliminate primer dimer and large size fragments, AMPure XP beads were then used to purify cDNA library. Then size distribution was analyzed on an Agilent 2100 bioanalyzer, a peak at around400~700bp range should be observed. Finally, the samples were ready for sequencing on Illumina Hiseq systems

### Cell sorting by FACS

Human CD34^+^ cells were suspended in 100μL of PBS+5% FBS for staining of antibodies. The following antibodies were used for cell sorting: CD34 PE (Biolegend, clone 581, cat #343506), CD34 FITC (BD bioscience, clone 581, cat #555821), CD38 PE-Cy7 (BD bioscience, clone HIT2, cat #560677), CD45RA APC/Fire 750 (Biolegend, clone HI100, cat #304151), CD49f BV421 (Biolegend, clone GoH3, cat #313623), CD74 PE (Biolegend, clone LN2, cat #326807), CD90 APC (BD bioscience, clone 5E10, cat #559869), CD114 PE (Biolegend, clone LMM741, cat #346105), CD115 PE (Biolegend, clone 9-4D2-1E4, cat #347303), CD71 PE (BD bioscience, clone M-A712, cat #561938), CD41 FITC (BD bioscience, clone HIP8, cat #561851), MHCii FITC (BD bioscience, clone Tu39, cat #562008). Cell sorting was performed on the BD FACSAria П.

### Colony forming assays

Colony forming assays for each population were performed by plating 1000 FACS sorted cells into MethoCult™ H4034 Optimum methylcellulose-based medium (StemCell Technologies). 3 independent experiments were performed for each population. After 14 days in culture, CFU-multilineage colonies were calculated according to instructions from StemCell Technologies.

### Single-cell qPCR

39 cells CD71^+^ clones, 38 cells from CD71^−^ clones, 39 cells from CD114^+^ clones and 39 cells from CD114^−^ clones were randomly picked from each clone. Individual primer sets (total of 96) were pooled to a final concentration of 0.1μM for each primer. Individual cells picked directly into 8-well strips loaded with 5μL RT-PCR master mix (2.5μL CellsDirect reaction mix, Invitrogen; 0.1μL RT/Taq enzyme, Invitrogen; 0.5μL primer pool; nuclease free water to 20 μL) in each well. Strips were immediately frozen on dry ice. After brief centrifugation at 4°C, Strips were placed immediately on PCR machine. Cell lyses and sequence-specific reverse transcription were performed at 50°C for 60 minutes. Then, reverse transcriptase inactivation and Taq polymerase activation was achieved by heating to 95°C for 3 min. Subsequently, in the same tube, cDNA was subjected to 20 cycles of sequence-specific amplification by denaturing at 95°C for 15sec, annealing and elongation at 60°C for 15 min. Pre-amplified products were diluted 5-fold prior to analysis. Amplified single cell samples were analyzed with Universal PCR Master Mix (Applied Biosystems), EvaGreen Binding Dye (Biotium) and individual qPCR primers using 96 × 96 Dynamic Arrays on a BioMark System (Fluidigm). Ct values were calculated using the BioMark Real-Time PCR Analysis software (Fluidigm).

